# Automated reconstruction of all gene histories in large bacterial pangenome datasets and search for co-evolved gene modules with Pantagruel

**DOI:** 10.1101/586495

**Authors:** Florent Lassalle, Philippe Veber, Elita Jauneikaite, Xavier Didelot

## Abstract

The availability of bacterial pangenome data grows exponentially, requiring efficient new methods of analysis. Currently popular approaches for the fast comparison of genomes have the drawback of not being based on explicit evolutionary models of diversification. Making sense of bacterial genome evolution, and notably in the accessory genome, requires however to take into account the complex processes by which the genomes evolve. Here we present the *Pantagruel* bioinformatic software pipeline, which enables the construction of a complete bacterial pangenome database geared towards the inference of gene evolution scenarios using gene tree/species tree reconciliation. *Pantagruel* is a modular pipeline that combines state-of-the-art external software with unique new methods. It can be executed with no supervision to perform a standard pangenome analysis, or be configured by advanced users to integrate methods of choice. A relational database underlies its data structure, allowing efficient retrieval of the large-scale data generated by integrative analyses of pangenome evolutionary history. From the reconstructed gene evolution scenarios, two main outputs are derived: firstly the gene tree-aware assignation of orthology, allowing the fine analysis of gene gain and loss history over the species phylogeny, and secondly a network of gene-to-gene association based on correlated events in scenarios of gene evolution, leading to the definition of co-evolved gene modules. *Pantagruel* is available as an open source software package at https://github.com/flass/pantagruel.

## Introduction

In recent years, increasing attention has been given to the study of bacterial pangenomes, in the hope of unravelling the genetic determinants of the complex phenotypic diversity of prokaryotic species (Vernikos et al. 2015). Such analysis is necessary to better understand how different strains are adapted to their environment – and even predict what are the set of environmental conditions in which they would strive.

However, the pattern of genomic gene content diversity is too complex to be deciphered at once: it is the result of a mixture of evolutionary processes by which genes propagate within and across genome lineages, including vertical descent (i.e. clonal replication of the gene along with the rest of the cell’s genome), homologous recombination of related gene alleles (between members of the same population or different but co-occurring populations (Didelot et al. 2010), and most importantly, horizontal gene transfer (HGT) – the event of introduction of a new gene (or allele) into a distant genetic background (Ochman et al. 2000). All these gene flow events occur at rates that are likely to vary from one cellular lineage to another, as well as between gene lineages.

As a result, the distribution of so-called accessory genes in strain genomes is neither clearly reflecting their vertical history of descent, nor completely dissociated from it (Konstantinidis & Tiedje 2005). This hinders efforts to identify lineage-specific genes involved in the adaptation of species to their ecological niche (Kumar et al. 2015), or conversely to pinpoint phenotype-associated genes using genome-wide association study approaches, where traits are supposed to be distributed independently of the non-causative genetic background (Earle et al. 2016; Collins & Didelot 2018). One consequence is that standard practice in microbial comparative genomics tends to ignore the accessory component of genomes, or to only consider the pattern of gene occurrence, disregarding the pattern of allelic variation, and the phylogenetic information it contains on the gene evolution.

There is therefore a need to develop a unifying framework for the study of microbial pangenome evolution. A central component is gene tree/species tree reconciliation methods (Szöllősi et al. 2015), that perform the comparison of the phylogenetic history of a gene, whether core or accessory, to the history of a reference tree. Here, the aim of the reference tree is not to reflect the clonal genealogy (Didelot et al. 2010) but rather to provide a common referential for the identification of events by which all genes propagate and diversify. Reconciliation methods thus deliver scenarios of gene family evolution, that depict the way genes evolved in and out of the frame of the reference tree. Once interpreted in the context of this unifying common frame, the phylogenetic tree of a gene is annotated with events of speciation, duplication, horizontal transfer or loss, that can for instance be compared with the history of mutational events along the same tree, thus allowing one to associate change in the gene sequence to events of dissemination across species.

This framework notably allows us to compare gene histories, and to identify segments of their past during which they co-evolved. This approach can document conserved associations between genes that go back much earlier than the most recent common ancestor of a clade of genomes that all contain these genes. For instance, the association between genes present on a plasmid carried by a particular clone most likely pre-dates its acquisition by the founder cell. Longterm co-evolution patterns can document selective pressures that constrained these associations, either because of epistatic interactions or through co-selection of genes in linkage. Co-occurrence patterns can similarly be interpreted as signatures of constrained co-segregation of genes in a population. However, the strength of association between physically linked genes can be inflated by recent events of selfish dissemination of a mobile genetic element (MGE) vector, or by strong but transitory selective regimes, like the selection of antibiotic resistance genes during antibiotic treatment. Recent co-acquisition of genes thus cannot be interpreted confidently as a sign of epistasis. By contrast, associations are unlikely to result from contingent causes if they were conserved over long periods of time during which purifying selection acted on the genomes. Moreover, genes that remain associated via events of co-transfer – which are unlikely under neutral conditions, due to random sampling of transferred genes and frequent genome rearrangements or gene losses – reflect the effect of strong selection for maintained gene linkage (Lassalle et al. 2017). This indicates that long-term co-conservation patterns are relevant signatures of selection that can be applied to study the evolution of accessory genes.

We introduce the bioinformatic software pipeline *Pantagruel*, a modular suite for the building of a complete bacterial pangenome database, including a reference species tree, homologous gene family alignments, gene trees and most importantly, gene evolution scenarios inferred using gene tree/species tree reconciliation. *Pantagruel* is implemented as a script that will deploy modules executing the various methods and third party-software as required. These modules can be used automatically in order for the integral building of a pangenome database, or independently by advanced users who desire to analyse intermediate output and/or to provide their own intermediary input files. Throughout the computations, results are gathered into a relational database which provides a naturally scalable interface for interconnecting the different aspects of the dataset and studying their patterns of association.

## Methods

The package *Pantagruel* provides an automated tool for the construction of a pangenome database from a large genome dataset and the application of phylogenetic reconciliation methods to all gene families in the pangenome. This is achieved through a parallelized workflow, which can be broken down into the ten main tasks described below.

### User interface

The *Pantagruel* interface relies on a single command-line executable `pantagruel` that can execute subprograms, or tasks, that are to be executed in order: `init` followed by tasks `0` to `9`. The `init` task sets up the parameters for the rest of the pipeline execution. Parameter values are passed on using command-line options and are stored in an environment file, a shell script defining environment variables that will be loaded at the beginning of every subsequent task. The environment file can be modified manually between tasks to adjust parameters, even though this is not recommended as it may create issues due to dependencies between tasks.

General usage is described using the `-h` option and in more detail on the code repository web page https://github.com/flass/pantagruel.

### Pipeline Tasks

A summary of the procedure used in each successive task of the pipeline is described below, while a detailed version is provided in the Supplementary Material. A schematic representation of the workflow is provided in Figure 1 (tasks init and 0-5) and Figure 2 (tasks 6-9).

**Figure 1:**
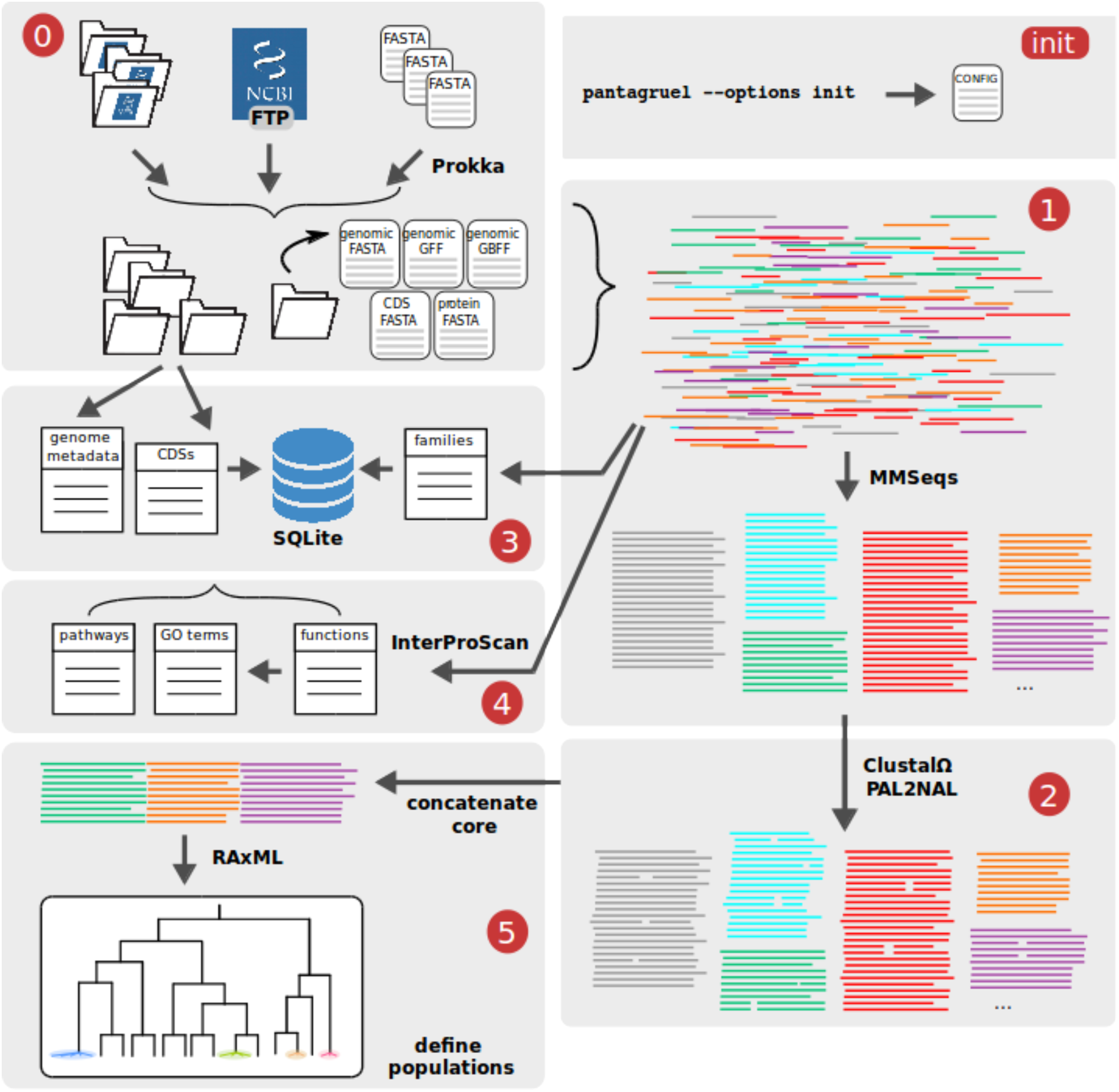
Schematic view of Pantagruel workflow for pipeline tasks init and 0 to 5. **(init)** user-defined parameters given via command-line options are stored into a configuration file that will direct the operations throughout the pipeline. **(0)** Input data are assembled from variable sources: annotated genome assemblies (formatted as) obtained from NCBI Assembly database; an assembly accession list for annotated genomes to be downloaded from NCBI Assembly FTP; or user-provided contigs of custom assemblies, which will be annotated by Prokka. **(1)** All proteins are extracted from annotated assemblies, and clustered into homologous families using MMSeqs. **(2)** Protein family sequences are aligned with ClustalOmega, and reverse-translated into CDS alignments with PAL2NAL. **(3)** A SQLite database is set up and loaded with data covering organism and genome assembly metadata, protein/CDS annotation and family clustering. **(4)** Proteins are uniformly annotated with functional domains using InterProScan, and these annotations are linked to Gene Ontology and metabolic pathway ontology terms; all annotation and ontologies are loaded into the SQLite database. **(5)** Core gene families are concatenated and used to infer a reference species tree with RAxML. Population of closely related genomes are delineated based on the tree topology.

**Figure 2:**
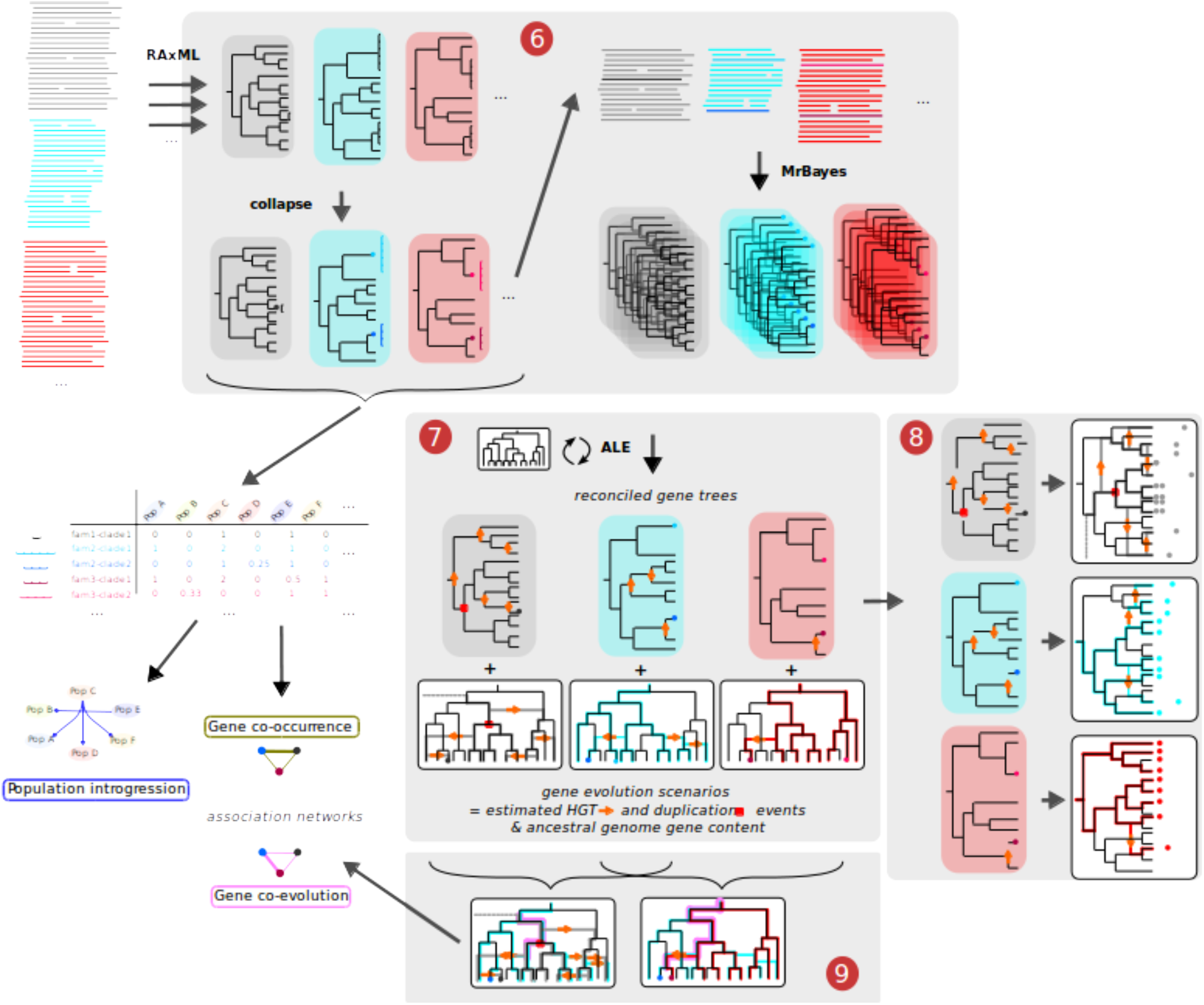
Schematic view of Pantagruel workflow for pipeline tasks 6 to 9. **(6)** From each gene family alignments, a rapid maximum-likelihood gene tree is estimated on the full alignment using RAxML. Clades of closely-related sequences with low topological supports within the subtree are collapsed. Alignments restricted to the sequence representative of collapsed clades (CCs) are used to perform a Bayesian sample of topologies of each gene tree using MyBayes. CCs also define tight clusters of similar sequences; the occurrence of CC cluster in the species populations (defined in task 5) is tabulated for further population genetics analyses. **(7)** Collapsed gene tree samples are reconciled with the reference species tree (from task 5) using ALE to infer reconciled gene trees annotated with events of horizontal transfer, duplication and loss of genes, as well as gene evolution scenarios placing these events in the reference species tree. **(8)** Based on the reconciled gene trees, groups of orthologs are defined within each homologous family, based on the gene tree topology and their common descent via events of speciation only. Phylogenetic profiles of occurrence of these ortholog groups are then used to infer sets of clade-specific genes for all clades in the reference tree. **(9)** Evolution scenarios are compared for all pairs of gene lineages, evaluating the extent of their co-evolution by exploring the reconciled gene tree from the gene tip to its root and summing the joint probability of common events. These coevolution scores are used to build a gene association network.

#### 0) Fetching of public genome assembly data and annotation of custom genome assemblies

Input genomic data can be provided in several ways: either as a list of NCBI Assembly accession numbers to be automatically downloaded from the NCBI FTP site, or directly as a collection of genome assembly files (Figure 1). The preferred format for assemblies is a full set of files as can be downloaded from NCBI Assembly database (RefSeq format), including genomic FASTA (contigs), genomic GFF (annotation), but also coding sequences (CDS) and protein FASTA files. Alternatively, input can simply consist of genome assembly contig files (FASTA), for instance when using newly sequenced genomes. Unless a custom genome annotation is provided in a specific format mimicking the NCBI Assembly format (see manual), genome annotation will be automatically made using Prokka (Seemann 2014). In the latter case, it is also possible to provide a custom set of genome assemblies to serve as references for a sequence similarity search leading the functional annotation of CDSs. Practically, annotation and sequence files analogous to the NCBI Assembly format are generated from the submitted contigs, including the extraction of all coding sequences (CDSs) and their translation into corresponding protein sequences.

#### 1) Building a homologous sequence database

All predicted protein-coding amino acid sequences are extracted from the genome assembly dataset and clustered into homologous gene families – which may include multiple copies per genome – using MMSeqs2 (Steinegger & Söding 2017).

#### 2) Aligning homologous sequences

Homologous amino acid sequences are aligned with ClustalOmega (Sievers et al. 2011), and codon alignments are generated by reverse-translating the protein alignments into their coding sequences using PAL2NAL (Suyama et al. 2006).

#### 3) Initiating SQL database

A relational database is set up, using a SQLite engine which is easily portable and avoids reliance on a client/server system requiring special administration rights. The database schema is designed to gather all data from further steps, and facilitate their exploration through SQL language database query.

#### 4) Functional annotation of proteins with biological ontologies

Proteins are systematically analysed using the battery of functional annotation tools gathered in InterProScan (Jones et al. 2014). InterPro domain annotation is translated into Gene Ontology (GO) terms and KEGG, BioCyc and Reactome metabolic pathway terms (Ashburner et al. 2000), providing unified frameworks for the functional analysis of further results.

#### 5) Estimating the reference tree

The reference tree is estimated from the concatenate of core genome gene alignments. The set of strictly core genes can be small when genome datasets are large and/or diverse, which would lead the derived tree to not be representative of the whole genome history. It is possible to relax the definition of this reference set to a pseudo-core gene set, defined as all gene families that are present in exactly one copy in almost all genomes in the dataset (e.g. 95%). Based on a concatenated alignment of these genes, a maximum-likelihood reference phylogeny is inferred using RAxML (Stamatakis 2014) under the GTRCATI model (25 site categories), and branch support are estimated based on parametric bootstraps. The tree root is inferred using the maximum ancestor deviation (MAD) technique (Tria et al. 2017). LSD (To et al. 2016) is used to generate an ultrametric version of the reference tree (dated in arbitrary units).

In order to correct for the bias in sampling of the phylogenetic diversity of the dataset and decrease the computational complexity associated with large phylogenies, we delineate clusters of very similar genomes, hereafter called populations, based on the (non-ultrametric) reference tree. We consider populations that can be nested, reflecting the recent emergence and evolution of bacterial clones from one another. Thus, while most often populations cover monophyletic sets of genomes, they are sometimes paraphyletic. The population delineation criterion relies on strong support of the stem branch and a long enough stem branch relative to the length of branches within the cluster; values of parameters can be modified through command line options.

#### 6) Estimating gene phylogenies

Gene diversification patterns result from several evolutionary processes, namely long-term diversification within genome lineages (subject to natural selection) and short-term dissemination within a meta-population formed of several genome lineages (likely subject to no selection or only transient episodes). The short-term diversification process likely involves phenomena in violation of the assumptions of phylogenetic methods (e.g. recurrent homologous recombination amongst highly similar sequences), and its common features with the long-term diversification process (e.g. occurrence of HGT) may present widely different parameter values (e.g. higher HGT rate observed over recent rather than ancient diversification; (Didelot et al. 2009, 2012)). Thus, the respective parts of gene sequence evolution that result from these two separate processes have to be identified, so that diversification can be modelled and analysed separately, under a phylogenetic framework and a population genetic framework, respectively.

Therefore, gene trees are estimated in several steps, with two objectives: (1) separating recent vs. ancient diversification for separate analysis; and (2) decreasing the complexity of data given as input to parameter-heavy phylogenetic inference methods.

In short, trees are first estimated with RAxML (Stamatakis 2014). Clades of gene sequences with low topological support (including groups of identical sequences) are iteratively collapsed, leaving only one representative leaf per clade. For each collapsed clade (CC), the frequency of each represented population (i.e. the count of leaves belonging to each population) is recorded for further population genetic modelling. A collapsed CDS alignment is generated to reflect the collapsed tree, and from this alignment a Bayesian sample of the topology of each ‘backbone’ gene trees is then obtained using MrBayes (Ronquist & Huelsenbeck 2003). Gene families are processed in parallel during this task, either using GNU Parallel (Tange 2011) when run on an isolated, multiple-core machine (e.g. a cloud server) or, when run on a computer cluster, by submission to worker nodes via a job scheduler (LSF and PBS submission systems (Altair 2019; IBM 2014; Adaptive Computing 2019) are currently supported).

With the objective of performing gene tree/species tree reconciliation, all leaves in the collapsed gene trees need to have a unique identity in the reference tree. A procedure is thus required to define which identity will be given to the leaf left to represent a CC. Closely related gene sequences occurring in genomes belonging to separate populations suggest they were recently exchanged by HGT between populations. In other words, these genes are assumed to be segregating in a meta-population formed of several populations, between which exchanges are possible despite their distant relationships. Under this assumption, it can be further considered that the gene was originally fixed in one ancestral population from which it diffused to other populations through HGT. We thus use a simple heuristic based on their occurrence profile to infer the population (as defined in task 5) where these sequences originated: the most densely represented population is the most likely ancestor. This inferred ancestral population is then used to relabel the leaf representing the CC. In certain cases, clusters of gene sequences belonging to monophyletic groups of populations suggest some sequences within the CC likely evolved vertically; in that case a subtree of population is grafted to the gene tree in place of the leaf representing the CC. These leaf relabelling or replacement operations are made on each tree of the Bayesian sample, thus preserving the diversity of topologies in the samples. Records of these operations are stored in the relational database initiated in task 3.

#### 7) Gene tree/species tree reconciliation

Dated gene tree/species tree reconciliation is conducted for each gene family to infer their scenarios of evolution, with events of gene duplication, transfer and loss (DTL) using ALE (Szöllősi, Tannier, et al. 2013). Gene DTL event rate parameters are set free and estimated by a maximum-likelihood (ML) approach, and then 1,000 gene family evolution scenarios and associated reconciliations are sampled using a Bayesian approach (Szöllősi, Rosikiewicz, et al. 2013). The output of ALE is then parsed to export inferred events to the relational database initiated in task 3, allowing to relate events involving CCs to their original constituent gene sequences.

#### 8) Inference of orthology from evolution scenarios

Using reconciliation data, we can define groups based on a formal criterion of orthology, i.e. common descent from an ancestor by means of speciation only (Doyon et al. 2011), rather than a proxy criterion such as bidirectional best hits (BBH) in a similarity search (Wolf & Koonin 2012; Dalquen & Dessimoz 2013). This means that any event of HGT or duplication annotated on a reconciled gene tree branch induces the subtree beneath to constitute a new orthology group (OG). However, this strict criterion tends to atomize families where most gene tree branches are annotated with transfer events even though each taxon is only represented once, for instance due to sparse taxonomic representation of the gene or to homologous recombination events. To account for this, we relaxed our OG definition and used a heuristic mixing the formal orthology criterion and the unicopy criterion (Bigot et al. 2013), where transfer events that do not increase the number of gene copies in a genome are ignored. We used a similar approach to differentiate additive and replacing gene transfer events in a previous study (Lassalle et al. 2017) where it provided an evolutionary sound, yet conservative way of describing the structure of diversity within gene families. A last hurdle in the definition of OGs from reconciled gene trees is that trees from a sample are not guaranteed to have the same topology, nor the same events associated with them. We therefore ran the orthology classification algorithm on each reconciliation in a sample. A network was then built that connects genes which were classified in the same OG in at least 50% of the sample; connected components of this graph provided the final OGs, as a consensus of the orthology structure inferred in every reconciliation of the sample.

#### 9) Quantification of gene co-evolution

The inferred event profiles of all genes in the pangenome are compared, quantifying their similarity with a score defined as the sum of joint event probabilities (SJEP). This score is obtained for a pair of gene lineages *g* and *h* by computing the sum for all events *e* of the probability that *e* affected both *g* and *h* (see Suppl. Material). The SJEP score thus describes the average number of evolutionary events in common during the history of two genes. Pairs of genes with a significant SJEP score resided together in ancestral genomes, notably for a number of ancestors on the reference tree at least equal to that score – possibly more if we consider that genes may be acquired in a same genome through independent events, for instance one by speciation and the other by transfer. Within the resulting matrix of pairwise SJEP scores, entries with high values were used to build a network of associations. The tightest hubs in this network reveal co-evolving gene modules, whereas connection between such hubs highlight recurring associations between modules, for instance between the large core genome gene module and smaller accessory gene modules.

The specificity of the SJEP score is that it describes a pairwise association between gene lineages, and can therefore detect localized co-evolution in a pair of gene trees, without the assumption that co-evolution must have involved the whole gene family. This allows a much more sensitive detection of gene associations, but introduce the problem of high-dimensionality of testing the association for the many combinations of gene lineages that can be enumerated within the dataset. For example, in a dataset of 1,000 genomes made of 5,000 genes, the number of pairwise comparisons is of the order of 10^13^. This computational challenge is addressed by an efficient use of relational database (SQL) queries in the search of significant matches between gene lineage evolutionary scenarios and by imposing filters on the probability of events that are supporting lineage matches (> 0.1), and on the co-evolution scores to report (> 1.0).

Even with these filters, the resulting network can be very dense. This potentially large number of association links is however redundant. In this gene lineage network representation, gene lineages from a same gene family are not independent as they share a significant fraction of their ancestral history. This may result in repeat association of closely related genes to the same set of genes. Redundant many-to-many relationships between two groups of close homologs can be simplified into a simple link between those groups.

For this reason, we aimed to group genes from a homologous gene family into sub-clusters. We define subgroups based on the orthology relationship (see task 8), as it ensures a fairly close relatedness between members and limits the size of a group to the size of the genome dataset. For each pair of OGs, we filter association links between member gene lineages to those with the highest score for each member gene lineage (best lineage hit) and report the mean of these best lineage hit scores as the OG-OG association score.

## Software implementation

*Pantagruel* is implemented as a master shell script commanding the task routines, written mostly in bash language. Each task script may itself call third-party programs or utility programs originally developed for *Pantagruel*-specific procedures, including phylogenetic tree manipulation and database query and dynamic management; these utility programs are mostly written in Python 2.7, as well as in R, bash and SQL languages. This script can easily be used to run the whole pipeline using a command-line interface. The full package is available on GitHub at https://github.com/flass/pantagruel.

*Pantagruel* was designed to run on a Linux server computer, and has been successfully tested on virtual machines 8 parallel cores, and 64 GB memory, as made available by the UK Medical Research Council Cloud Infrastructure for Microbial Bioinformatics (MRC CLIMB) (Connor et al. 2016).

A script is also provided to automatically install the dependencies. This script is fully compatible with Debian-type Linux operating systems such as Ubuntu; it may also be used on other Linux or Unix systems but some manual steps may be required for complete installation.

Computationally intensive pipeline tasks such as gene tree estimation and gene tree/species tree reconciliation (tasks 6 and 7, respectively) spawn many independent computation jobs (one per gene family) that naturally lend themselves to parallelization. For these tasks, specific scripts are provided to execute them on a high-performance computer (HPC) cluster to speed-up the execution; only PBS/Torque job submission systems are supported so far.

In addition, another version of the package is in development using the Bistro framework (https://github.com/pveber/bistro), to provide a pre-compiled, self-contained Docker image that can be deployed on pretty much any platform, including swarms of virtual machines (VMs).

## Discussion

*Pantagruel* is a comprehensive pipeline for the evolutionary analysis of bacterial pangenome datasets. A central component is the use of reconciliation methods (Szöllősi et al. 2015) to compare the phylogeny of a given gene with that of the genome as a whole. State-of-the-art reconciliation techniques are however too computationally expensive to be applied to hundreds of genomes each containing thousands of genes. We therefore implemented within *Pantagruel* new ways to simplify the input data to ease the computation of reconciliations with limited loss of information. Clades that cluster very similar gene sequences and show little internal topological support are separated from the well supported part of the gene tree (Figure 2). The absence of phylogenetic information within those clades, and the possibility of homologous recombination having occurred between such closely related sequences, justify that their relationships should not be considered under a phylogenetic model. Instead, the closely related sequence clusters indicate recent gene sharing, where population genetics and ecological modelling approaches would be more appropriate. Notably, combining the information of population distribution in all such closely related sequence clusters into a network would allow to identify meta-populations within which closely related populations preferentially exchange genes (Ansari & Didelot 2014). We also use this information to infer the identity of the recent ancestors at the stem of these rake clades, which can be annotated on the tree backbone and fed back to the reconciliation method. Thanks to this separation of data, the phylogenetic reconciliation inference problem is largely simplified and solutions concerning the well resolved parts of gene phylogenies are found to be more precise. We observed that this way, the reconciliation software achieved large speed-ups, but also could simply be run on gene families in which the initial complexity otherwise led to systematic excess memory use and termination. *Pantagruel* thus allows the practical use of reconciliation methods (we used ALE but in principle other methods could be used) on large-scale pangenome datasets.

Methods for statistical testing of gene lineage association will be developed using simulation-based approaches as previously used for phylogenetically-correct testing of phenotype-genotype associations in clonal microbes (Collins & Didelot 2018). This will be based on simulations of genome evolution under a similar DTL model that accounts for gene linkage evolution. Simulation could be provided by pangenome tree simulators such as FwdTreeSim (https://github.com/flass/FwdTreeSim) or Zombi (Davin et al. 2018). Development of analytical solutions to evaluate the likelihood of observing co-evolving pairs under a neutral model is also envisaged, notably using a linear algebra framework (Behdenna et al. 2016).

## Author contributions

FL and XD designed the method, FL wrote the original software and implemented the script version of the pipeline, PV implemented the Bistro/Docker version of the pipeline, FL and EJ tested the software. FL and XD wrote the manuscript. All authors have read and agreed to the content of the manuscript.

## Acknowledgements

We thank Eric Tannier, Gergely Szöllősi, Vincent Daubin and Rafal Mostowy for fruitful discussions about developing the algorithm for tree collapsing, integration of ALE and general pipeline feature development. Test computational calculations were performed with Imperial College HPC and MRC CLIMB cloud-based computing servers. This work was supported by Medical Research Council (MRC) grant MR/N010760/1.

