## Supplementary Material for "Automated reconstruction of all gene histories in large bacterial pangenome datasets and search for co-evolved gene modules with Pantagruel"

*Supplementary Material: Detailed description of pipeline procedure.*

### *Pantagruel task* **`init`**

At this step, options are processed to set up parameters. Mandatory options include -d and -r, which define the name and the location of the database to be created, as well as one or a combination of -a, -A or -L, which specify the input genome assembly dataset. Other options are facultative and their omission results in adopting default values for runtime parameters.

Parameters can only be passed on using command-line options during the `init` task, during which they are stored in the database environment file. The environment file can be modified manually between tasks to adjust parameters, even though this is not recommended as it may create issues due to dependencies between tasks.

### *Pantagruel task* **`0|00|fetch|fetch_data`**

In this task, *Pantagruel* will process the input genome data and verify its format validity. Note that for the sake of computing evolutionary analyses that have any meaning at all, *Pantagruel* requires that you provide a minimum of four genomes in input – ideally much more, as *Pantagruel* can easily deal with several hundreds of genomes. This minimum number 4 is to be split at the user's discretion between NCBI Assembly-type and custom assemblies, through `-A`,`-L` and `-a` options, respectively.

Custom genomes sequences lacking annotation files are annotated with Prokka (version 1.12) (Seemann 2014), with the following notable options: `--force --addgenes --locustag --compliant --usegenus --genus ${refgenus} --kingdom Bacteria --gcode 11`. `${refgenus}` is a variable which indicate a subject database for a BLASTP (Camacho et al. 2009) similarity search during the Prokka annotation process. Its value can be edited manually in the environment file to indicate a specific BLASTP database to Prokka; this database needs to be present in the db/genus/ folder of your Prokka installation (default contains reference databases for *Enterococcus*, *Escherichia* and *Staphylococcus*). By default, ${refgenus} value is 'Reference', which triggers the building of a reference BLASTP database from the combined proteomes of input genome assemblies that are already annotated, specifically those provided in the NCBI Assembly format through options `-A`,`-L`, `--refseq_ass4annot` or `--refseq_list4annot`. These proteomes are merged into a non-redundant dataset that is then thinned by retaining a single representative protein for each protein cluster as determined by the Linclust clustering program (in MMseqs package version 10-6d92c) (Steinegger & Söding 2018), using the `easy-linclust` command with default parameters.

### *Pantagruel task* **`1|01|homologous|homologous_seq_families`**

Proteomes from all input genome assemblies are merged into a single dataset. MMseqs2 release 2 (Steinegger & Söding 2017) is used (with default parameter values) to search for similarities between all pairs of proteins and to cluster them into families of homologs (not orthologs: homology relationship is inferred irrespective of the occurrence pattern in genomes). A first search is conducted by the clusthash module with 100% sequence identity criterion to dereplicate the sequence database. Then, sequence similarity search and clustering are conducted under default settings on the non-redundant protein sequence set.

***Pantagruel task* `2|02|align|align_homologous_seq`**

Protein sequences from each homologous family are aligned using ClustalOmega (Sievers et al. 2011) and reverse-translated into coding sequence (CDS) alignments using PAL2NAL (Suyama et al. 2006). Parallelization of alignment and reverse-translation tasks is done with GNU parallel (Tange 2011) and a with Python wrapper script using the `subprocess` module, respectively.

***Pantagruel task* `3|03|sqldb|create_sqlite_db`**

A relational SQL database is created with an SQLite engine. Its schema is designed to gather all data from further steps. Genome assembly and gene annotation data, as well as homologous gene classification are loaded in the database at this step.

***Pantagruel task* `4|04|functional|functional_annotations`**

In addition to Prokka annotation, functional annotation of proteins is done using the standalone InterProScan tool (Jones et al. 2014) with options '`--iprlookup --goterms --pathways`' to add annotation of InterPro terms, Gene Ontology (GO) terms and KEGG, BioCyc and Reactome metabolic pathways, providing an annotation with controlled vocabulary that is homogeneous across the dataset. When installing *Pantagruel,* the last version of InterProScan (currently 5.33-72.0) is automatically fetched, which enables the look-up of precomputed annotation in the current InterProScan database release and allows for a significant speed-up.

***Pantagruel task* `5|05|core|core_genome_ref_tree`**

a) Estimation of the reference phylogeny from pseudo-core gene concatenated alignment

The set of pseudo-core gene families is defined as all gene families that are present in exactly one copy in at least *p* genomes in the dataset. Default behaviour is to use *p* = *N* the total number of genomes, i.e. to use a strict core genome definition; this is however not advised, as the resulting genome set can be very small and yield a bad representation of the genome history. No given formula can provide the optimal set for every dataset because the structure of diversity cannot be predicted – not before running the program; notably, genome sequences with reduced gene content or partial sequences resulting from low-depth sequencing may lead to a biased distribution of missing genes (absence of genes always in the same genome) that could later distort the phylogenetic inference. It is therefore advised to use the interactive script '`choose_min_genome_occurrence_pseudocore_genes.sh`' that is provided with the *Pantagruel* package to explore the distribution of pseudo-core gene presence and absence across genomes. Based on a concatenated alignment of these genes, a maximum-likelihood reference phylogeny is inferred using RAxML (version 8.2.11) under the GTRCATI model (25 site categories), and branch support are estimated based on parametric bootstraps. The tree root is inferred using the maximum ancestor deviation (MAD) technique (Tria et al. 2017). LSD (To et al. 2016) is used to generate an ultrametric version of the reference tree (dated in arbitrary units) that will be later used for reconciliation with ALE.

b) Identification of genetically monomorphic populations from the reference tree

Populations are defined by exploring the tree, following a pre-order traversal (from tips to root), and performing an iterative pruning of groups recognized as populations. The procedure is the following: given a subtree $t_i$ has a stem (root-side) branch of length ≥ $l_s$ and support ≥ $b_s$ and with all internal (within the subtree) branches of length < $l_w$, a population $p_i$ of $n_i$ members corresponding to the leaves of the subtree $t_i$ is registered. In the case of a genome with no close relative, a trivial population with a single member is registered. After registering a population $p_i$, the population subtree $t_i$ is masked for the rest or the tree traversal, allowing the definition of another population $p_j$ from a subtree $t_j$ that would encompass $t_i$.

### *Pantagruel task* `6|06|genetrees|gene_trees`

In order to mask unresolved parts of the gene tree topology that may confuse later phylogenetic inferences and to reduce the dimension of the gene trees prior to compute-intensive gene tree/species tree reconciliation inference with ALE (Szöllősi, Tannier, et al. 2013), a data reduction approach is implemented. It is similar to that described in (Izquierdo-Carrasco et al. 2011) in that it aims at recognizing the backbone of a gene tree topology, but differs in its main purpose, which in our case is to discard parts of the tree that are not well supported, and only as a consequence to reduce the required computation time and space. This drove us to implement different criteria for the recognition of subtrees to prune off the tree backbone, which are based on topological support instead of the set reduction factor used by Izquierdo-Carrasco et al. 2011.

a) First estimation using maximum-likelihood on full dataset

For all gene families with at least four sequences, a starting quick ML estimate of each gene tree *G* is obtained from the full CDS alignments using RAxML (version 8.2.11) under the GTRCAT model (25 site categories), and branch supports are estimated based on 100 rapid bootstraps.

b) Collapsing low-diversity clades

Clades of gene sequences with low topological support are identified for collapsing. Clades with strong stem branch support (default: ≥ 70% branch support) but low support for their subtree topology (default: median branch support ≤ 35% and maximum branch support < 70%) are collapsed, i.e. all sequences in the clade are removed from the alignment, apart from one random sequence that is kept as a representative of the clade diversity; this representative is given an arbitrary label 'clade$_n$' (with *n* an integer) with no species identity. This results in an aligned sequence dataset representative of the backbone of the phylogenetic diversity of the gene family. In addition, for each collapsed clade (CC), the frequency of each represented population (i.e. the count of leaves belonging to its member species) is recorded.

With the objective of performing gene tree/species tree reconciliation, it is crucial that all leaves in the collapsed gene trees are given a species identity. A procedure is thus required to define which identity will be given to the leaf left as representative of a CC. Closely related gene sequences occurring in genomes belonging to distant populations suggest they were recently exchanged by horizontal transfer between populations. In other words, these genes are assumed to be segregating in a meta-population formed of several distantly-related populations, between which exchanges are possible irrespectively of their relationships. Under this assumption, it can be further considered that the gene was originally fixed in one ancestral population from which it diffused to other populations through random mating. It is thus possible to infer the population of origin of these sequences using a simple heuristic based on their occurrence profile: the most densely represented population is the most likely ancestor.

However, it is also possible that some genes that have been vertically transmitted over long times have so strongly conserved sequences that they form clades matching the criterion of high sequence similarity for identification of CCs. We thus aim to distinguish clades of ancient, vertically transmitted sequences from clades of recent horizontally disseminated ones. We first regroup populations represented in the CC according to their distribution in $R_c$: we group together populations forming monophyletic groups, and aggregate the monophyletic groups which ancestors are connected by a path of inter-node length ≤ 4. This threshold allowed the best recovery of groups of clonal inheritance pre-dating the advent of species populations as defined above, while accounting for potential losses in nested lineages. Having identified populations or groups of populations in which the genes were likely clonally inherited, we rank the putative ancestral (groups of) populations $p_i$ following this series of criteria:

- having a median frequency of the gene f($p_i$) > 0.5 (boolean meaning that at least half the members possess the sequence) – this binary criterion aims at discarding populations with low occurrence without giving too much importance to the observed frequency, which can be highly biased by the sample size;
- older age of the population (group) ancestor in the dated reference tree – this is to give priority to the oldest group as the progenitor of the gene lineage;
- higher frequency of the gene f($p_i$) – to break ties among contemporaneous populations.

The best ranking population is selected as putative origin in the recent dissemination process, and is subsequently used to annotate the species identity of the CC (see subsection 4d below).

This heuristic could be replaced by other methods, including methods based on a population genetics framework, or methods of ancestral state reconstruction retracing the history of genes from their species distribution (phylogenetic profile) and accounting explicitly for the species tree history. Integration in *Pantagruel* of the ancestral state reconstruction method Count (Csűrös 2008; Csűrös & Miklós 2009) for this purpose is considered, as it accounts for the possibility of vertical evolution resulting in clusters of gene sequences from monophyletic groups of populations.

c) Bayesian estimation of backbone gene trees

A Bayesian sample of the topology of each 'backbone' gene trees {$G_b$} is obtained from the collapsed CDS alignment using MrBayes (version 3.2.6) (Ronquist & Huelsenbeck 2003) under the GTR model. Two independent Metropolis-coupled Monte-Carlo Markov chains (MCMCMC) runs are carried out over 2,000,000 iterations of 4 coupled chain each, sampling a tree from the cold chain every 500 for output, resulting in 2 times 4,000 sampled trees. At the end, samples are compared between runs using the sumt command to produce convergence diagnostics (excluding the first 25% of the sample as burn-in); if convergence is not achieved (based on the criterion of standard deviation of split frequencies > 0.10), another round of 2,000,000 generation is run. This is repeated until reaching convergence, or after 5 rounds.

Gene tree samples from different families are computed in parallel. When run on a single isolated machine (with multiple nodes), e.g. a cloud server, MrBayes is run on a single core per family and parallelism between families is managed using GNU Parallel (Tange 2011). When run on a computer cluster, MrBayes is run on 8-core worker nodes and jobs are submitted via a job scheduler (LSF and PBS submission systems (IBM 2014; Altair 2019; Adaptive Computing 2019) are currently supported).

d) Joint collapsing of species and gene trees

In each tree from the Bayesian sample, anonymous tips representing CCs are re-labelled with their inferred ancestral population identity (see above). If the origin of the CC genes is inferred in a single population, the anonymous leaf is simply labelled with this population name. If the CC origin is inferred in a group of related populations, a replacement clade (RC) depicting the population relationships (i.e. with population-labelled leaves) is grafted to the gene tree. To do so, the subtree descending from the last common ancestor of the population group is extracted from $R_c$ and is pruned so to only feature the represented populations. Then, the branches of the subtree are scaled to match the relative depth of the original CC in the full ML gene tree. Finally, the obtained RC is substituted to the anonymous tip representing the CC. This procedure aims at best representing the occurrence of the genes in the population of origin (accounting for losses), while discounting the result of recent horizontal spread. This way, the local reconciliation of the grafted subtree with the reference tree yields a locally parsimonious scenario of evolution (only involving speciation and loss events) that will not impact the reconciliation of the rest of the gene tree. We thus obtain a collapsed gene tree sample $\{G_c\}$ with consistent tip labelling, but variable topologies and branch lengths – apart from the grafted subtrees that are homogeneous among the sample.

Finally, we take care of restoring the matching of the leaf labelling between the gene trees and reference species population tree $R$. For each gene family $g$, a family-specific reference tree $R_c(G)$ is generated. In order to minimize the size of the resulting reference tree and so to decrease the complexity of following reconciliation computations, we set to collapse the maximum of populations as in $R_c$, while preserving the reference-to-gene tree species label matching. We consider the following species categories for each population $p$: $\{m\}$, the set of member species of $p$ represented in $\{G\}$ can be divided into three sets: $\{m_{\mathrm{CC}}\}$, the set of member species only represented within CCs of $\{G\}$, $\{m_{\mathrm{lone}}\}$ the set of member species represented only outside CCs of $\{G\}$, and $\{m_{\mathrm{both}}\}$, the set of member species occurring both within and outside of CCs. We proceed as follows:

i) When $\{m_{\mathrm{lone}}\}$ set is null, the population $p$ is collapsed in $R_c(G)$, i.e. replaced by a $p$-labelled tip.

ii) When $\{m_{\mathrm{lone}}\}$ set is not null, the species in $\{m_{\mathrm{lone}}\}$ are made to feature in $R_c(G)$, while the remainder of $p$ is represented by an additional collapsed $p$-labelled population tip. The $p$ and $\{m_{\mathrm{lone}}\}$ labels then form a monophyletic group in $R_c(G)$ and their relationship follows those in $R$.

iii) When $\{m_{\mathrm{lone}}\}$ set is not null and $|\{m_{\mathrm{CC}}\} \cup \{m_{\mathrm{both}}\}| = 1$, the species in $\{m_{\mathrm{lone}}\}$ are made to feature in $R_c(G)$, while the single species represented in a CC is represented as itself, i.e. all the species in $\{m\}$ are represented in $R_c(G)$ as in $R$.

In cases i) and ii), if the set $\{m_{\mathrm{both}}\}$ is not null, another normalization operation needs to be done in the gene tree sample $\{G_c\}$, by converting the species labels of the lone tips (i.e. those occurring outside CCs) into the label of their population. While this adjustment is necessary to keep the bijective relationship between species and gene tree labels, it could be considered artificial and possibly introduce artefacts in reconciliation estimation. However, we consider this operation mostly safe due to the relative low frequency of this type of adjustment. In addition, we consider the fact that those lone tips most often occur in the gene trees far from representatives of their closely related species, and thus would most likely be recognized as recipient of transfer events, regardless of their identity as a single species or population, thus not changing the overall outcome of the gene evolution scenario.

e) population of relational database with phylogenetic information

As a final step, the relational database set up during *Pantagruel* task 3 will be populated with data describing the collapsing of clades in initial gene trees and their replacement by representative tips of subtrees. This allows to rigorously keep track of the correspondence between actual sequence and their representation in reconciled gene trees in the following tasks.

***Pantagruel task*** `7|07|reconciliations`

We use gene tree/species tree reconciliation software ALE (version 0.4) (Szöllősi et al. 2012; Szöllősi, Tannier, et al. 2013; Szöllősi, Rosikiewicz, et al. 2013) to estimate gene evolution scenarios under a model with gene duplication, transfer and loss (DTL). We use as input the pairs of matching reference trees $R_c(G)$ and gene tree samples $\{G_c\}$. In the latter, we retain the last 6,000 sampled trees, corresponding to 25% burn-in fraction in most case, or 50% for chains that were sampled for an extra round of MrBayes estimation. The ALE software has several modes, including the ALEml_undated and ALEml algorithms that implement the undated and dated xODT models, respectively. We apply the dated ALEml algorithm that considers time contraints in horizontal transfer, i.e. that a gene transfer cannot occur between species tree branches that do not overlap in time. ALE first estimates the event rate parameters of the DTL model using a maximum-likelihood approach and then samples of 1,000 scenarios from the reconciliation space using a MCMC walk. The output of ALE is parsed to export inferred events to a database. Each reconciled gene tree in the sample is processed as follows:

i) the list of atomic events $e$ is identified on each branch and recorded as ($X$, $b$) recording their type $X \in \{D, T, L, O, S\}$ (for duplication, transfer, loss, origination, speciation) and their location on branch $b$ of reference tree $R$. In the special case of transfers, the location of both donor and recipient species are recorded as (T, $b_d$, $b_r$). Events associated with branches from grafted RC subtrees are artificial and are therefore discarded.

ii) For efficient storage of the information, all events from all gene families are recorded in reference to a database listing all possible events. To allow later comparison of scenarios across families, events are recorded using the same referential of events as enumerated on the collapsed reference tree $R_c$. Because of the difference of leaf set between each $R_c(G)$ and $R_c$, events estimated by ALE with $R_c(G)$ as reference are translated into the referential of $R_c$. This is straightforward as every $R_c(G)$ instance shares with $R_c$ the branches leading to the population ancestors. We thus relocate the occurrence (or reception for T events) of every event inferred on branches of $R_c(G)$ within the population clade to the last common ancestor of that population. This procedure results in the truncation of gene history. Later events, i.e. L or S events inferred on branches located within population clades, are instead recorded in the same framework than the segregating presence/absence state of CC genes among populations (see section 5b above).

iii) because the same event located in the reference tree can occur in distant part of a gene tree (e.g. similar transfer event occurring in diverged clades of paralogs), we record the gene lineage, i.e. the tip-to-root path, on which the events are located. We thus obtain an ordered list of events $L(g, G_i)$ for each gene sequence $g$ in each reconciled gene tree $G_i$. To integrate results over the reconciliation sample, we estimate the probability $P(e|g)$ of having the event $e$ in the ancestry of gene $g$ as the frequency at which the event $e$ was inferred above the gene tip $g$ in the sample of all gene trees $G_i$ :

$$P(e|g) = \Sigma_i x_i \text{ where } x_i = 1 \text{ if } e \in L(g, G_i) \text{ and 0 otherwise.} \quad (1)$$

$P(e|g)$ is recorded for every observed combination of $g$ and $e$. This method of storage does not take into account the topology of the gene tree and is therefore highly redundant, notably regarding the ancient history of related gene sequences. However, this remains the only correct way without making any assumption on the topology and root of the gene tree, which can vary widely between sampled reconciliations due to the aggregation of topologies from the Bayesian gene tree sample (Szöllősi, Rosikiewicz, et al. 2013). The indexing of $P(e|g)$ by key tuple ($e$, $g$) in a relational database framework is used to allow their efficient query in following steps.

***Pantagruel task*** **`8|08|specific|clade_specific_genes`**

Based on the estimated gene family evolution scenarios, we could define gene groups based on a true criterion of orthology, i.e. common descent from an ancestor by means of speciation only (Doyon et al. 2011), rather than a proxy criterion such as bidirectional best hits (BBH) in a similarity search. This has the advantage of explicitly detecting the gain of an orthologous group (OG) in a genome lineage by means of horizontal gene transfer (HGT) or gene duplication. To differentiate additive HGT and replacing HGT events (i.e. gene conversion by homologous recombination events), we used a heuristic based on the unicopy criterion (Bigot et al. 2013) as described in a previous study (Lassalle et al. 2017), where transfer events that do not increase the gene copy number in a genome are not interpreted as the emergence of a new OG, i.e. considering homologous recombination can occur within an OG without breaking it. This OG classification heuristic was applied to each sampled scenario among the 1,000 trees sampled for each gene family. To summarize these data, a network was built that connects genes classified in the same OG in more than 50% of the sample; connected components of this graph provided the final consensus OGs.

This classification is then used to build a matrix of OG presence/absence in the dataset, and to compute the set of genes specifically present or absent in each clade of the species tree. Clade-specific presence (absence) is defined under the criterion that every OG present (absent) in all members of the focal clade and absent (present) in all members of its sister clade. It was previously shown that relaxing this criterion by allowing 'leaky' presence/absence in the contrasting species group to allow for additional event of transfer or loss led to better capture functional groups of genes in clade-specific sets (Lassalle et al. 2011, 2017). The default strictness of this criterion (zero presence/absence in contrasting species) can thus be relaxed by editing the file ending by '_clade_defs' in the folder '05.core_genome/' of the database; It is also possible to define custom species sets – that do not have to form clades in the species tree – for specific gene content comparison.

***Pantagruel task*** **`9|09|coevolution`**

The inferred event profiles of all genes in the pangenome are compared, quantifying their similarity with a score defined as the sum of joint event probabilities (SJEP). This score is obtained for a pair of gene lineages $g$ and $h$ by computing the sum of joint probabilities of events inferred on both lineages $P(e|g,h) = P(e|g) \cdot P(e|h)$ for all events $e$:

$$\mathrm{SJEP}(g, h) = \sum_e P(e|g,h) = \sum_e P(e|g)\, P(e|h) \qquad (2)$$

The SJEP score thus describes the average number of evolutionary events in common during the history of two genes. This implies these genes resided together in ancestral genomes, notably for a number of ancestors located at the species tree nodes at least equal to that score – possibly more if we consider that genes may be acquired in a same genome through independent events, for instance one by speciation, the other by transfer. The resulting matrix of pairwise SJEP scores, entries with top association values were used to build a network of associations. The tightest hubs in this network reveal co-evolving gene modules, whereas connection between such hubs highlight recurring associations between modules, for instance between the large core genome gene module and smaller accessory gene modules.

The specificity of the SJEP score is that it describes a pairwise association between gene lineages, and can therefore detect localized co-evolution in a pair of gene trees, without the assumption that co-evolution must have involved the whole gene family. This allows a much more sensitive detection of gene association, but introduce the problem of high-dimensionality of testing the association for the many combinations of gene lineages that can be enumerated within the dataset – in a dataset of 1,000 genomes made of 5,000 genes, the number of pairwise comparisons that can be done is of the order of $10^{12}$. This computational challenge is addressed by an efficient use of relational database (SQL) queries in the search of significant matches between gene lineage evolutionary scenarios and by imposing filters on the probability of events that are supporting lineage matches (> 0.1), and on the co-evolution scores to report (> 1.0).

Even applying this filter, the resulting network can be very dense. This potentially large number of association links is however redundant. In this gene lineage network representation, gene lineages from a same gene family are not independent as they share a significant fraction of their ancestral history. This may result in repeat association of closely related genes to the same set of genes. Redundant many-to-many relationships between two groups of close homologs can be simplified into a simple link between those groups.

For this reason, we aimed to groups genes from a homologous gene family into sub-clusters. We define subgroups based on the orthology relationship (see task 8), as it ensures a fairly close relatedness between members and limits the size of a group to the size of the genome dataset. For each pair of OGs, we filter association links between member gene lineages to those with the highest score for each member gene lineage (best lineage hit) and report the mean of best lineage hit scores as the OG-OG association score.
